# Reading papers: Extraction of molecular interaction networks with large language models

**DOI:** 10.1101/2025.07.21.665999

**Authors:** Enio Gjerga, Philipp Wiesenbach, Christoph Dieterich

## Abstract

**Motivation:** Signalling occurs within and across cells and orchestrates essential cellular processes in complex tissues. Cell signalling involves several different components, including protein-protein interactions (PPI) and transcription factors (TF), to promoter binding in gene regulatory networks (GRNs). Dynamically changing conditions oftentimes lead to the rewiring of cellular communication networks. Computational modelling approaches typically rely on databases of possible molecular interactions. Evidently, manual curation of databases is time-consuming and automatic relation extraction from scientific literature would greatly support our strive to understand molecular mechanisms. To ease this process, we reason that prompt-based data mining with Large Language Models (LLMs) could be used to extract information from relevant scientific publications.

**Approach:** In our work, we use open-source LLMs to mine an annotated corpus of molecular interactions. We focus on the extraction of entity relations between proteins, as exemplified in protein-protein interaction networks, and transcription factor to target gene relations, as exemplified in gene regulatory networks.

**Results:** We obtain promising evaluation results as measured by precision, recall and F1-score for the extraction of PPI relations: 87%, 70% and 71% and 77%, 57% and 62% for GRN relation extraction over a large corpus of short (average 331 tokens) scientific texts.

**Availability:** Codes with scripts and results have been provided in: https://github.com/dieterich-lab/LLM_Relations.

## 1. Introduction

Understanding molecular cell and tissue biology requires a comprehension of how biological molecules such as proteins, DNA and RNA interact with each other in space and time. These dynamic molecular interactions regulate important cellular processes such as growth, division, repair, response to pathogens, etc. Depending on the nature of the relation, molecular interaction networks can be categorised into different types. For example, protein-protein interactions (PPIs) are responsible for the regulation of fundamental processes such as signal transduction and the regulation of enzymatic activities. Gene regulatory networks (GRNs), on the other hand, control the expression of specific genes, thus defining cellular states and their fate. Any dysregulation of these interactions may cause diseases, including cancer, autoimmune disorders, cardiac diseases, etc.

Prior knowledge of interactions and relations involved in cell signalling can serve as a basis for the functional and mechanistic study of pathway systems (Aldridge et al. 2006). Databases of prior knowledge have been established throughout the years, and they provide resources for interaction and relations between signalling molecules, such as Protein-protein interactions (PPIs) (Szklarczyk et al. 2022; Oughtred et al. 2021; Del Toro et al. 2022; Türei et al. 2021; Xenarios et al. 2000; Albrecht et al. 2025) and GRNs (Garcia-Alonso et al. 2019; Liska et al. 2022). However, these resources typically rely on manual curation, which is prone to bias as well as human error. Additionally, keeping databases of molecular interactions up-to-date is a daunting task with the ever-growing amount of published literature. Given this, the automated and efficient extraction of molecular interaction relationships has the potential to significantly improve the speed and precision of creating knowledge resources that are domain-specific to a particular field, disease, or condition and aligned with the latest literature.

In this aspect, the emergence of LLMs has unlocked new opportunities for mining domain-specific molecular interactions from literature. These models, while not necessarily exclusively trained on scientific literature, can be used to readily extract knowledge in the form of entity relations for any type of scientific domain (Zhao et al. 2023). However, despite the excitement, prompting LLMs may be prone to introducing considerable errors, including hallucinations (interactions between absent entities), false negatives (missing out on true interactions), and false positives (reporting non-existent or incorrect interactions of present entities). Considering this, a rigorous evaluation of LLMs relation extraction (RE) capabilities in the context of molecular interactions would be of great relevance prior to widespread adoption. Our study focuses on prompt-based RE and uses open-source LLaMA-v3.1 with 8b parameters (from hereon referred to as Llama-8b) as well as LLaMA-v3.3 with 70b parameters (from hereon referred to as Llama-70b) LLM for PPI and GRN REs.

## 2. Methods

### 2.1. Relation Extraction Workflow

We have established an RE workflow that utilises prompt-based data mining of LLMs to obtain information on two crucial aspects of cell signalling: PPIs and GRNs (Figure 1). The workflow takes as input a collection of scientific text or literature which, when presented in a PDF format, needs to be converted into a structured format through parsing. For the extraction procedure itself, we employ open-source LLMs, namely LLaMA-v3.1 with 8B parameters (Llama-8b) and the larger LLaMA-v3.3 with 70B parameters (Llama-70b) models (Touvron et al. 2023); (Grattafiori et al. 2024).

**Figure 1.**
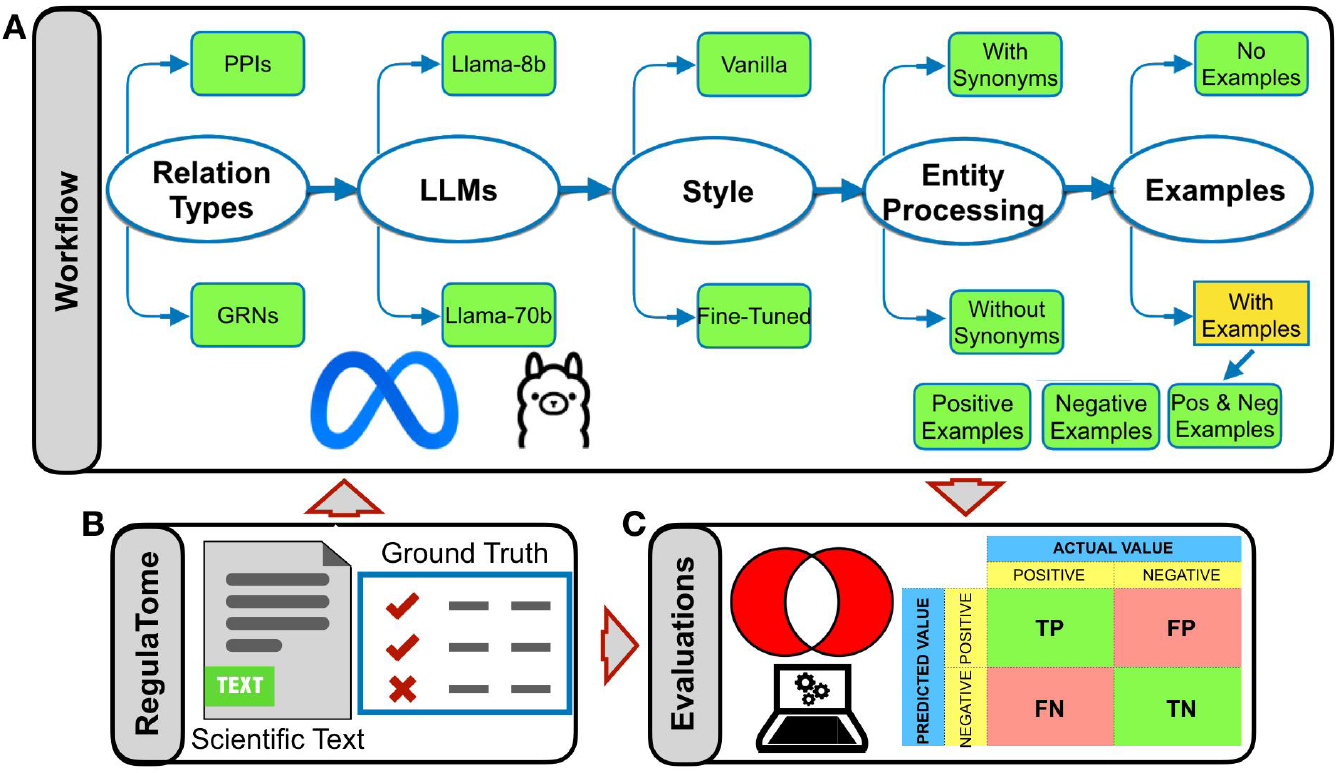
The Relation extraction workflow. **A.** The workflow involves the extraction of PPI and GRN relation types by using Llama-8b and Llama-70b LLMs in either the Base pre-trained (Vanilla) or Fine-tuned mode, as well as by supplementing the prompts with external examples of how a relation should and/or should not be extracted, as well as without providing any such examples. Before evaluation, the entities of the extracted relations can either be assigned or not, with up to 10 synonyms from the LLMs. **B**. The annotated RegulaTome corpus was used to extract the PPI and GRN relations. **C**. Evaluations of the extractions were performed in terms of precision, recall and F1-score values.

Both Llama-8b and Llama-70b language models represent auto-regressive language models built on an optimised transformer architecture. Their fine-tuned versions leverage supervised fine-tuning (SFT) and reinforcement learning with human feedback (RLHF) to align the model’s outputs with human preferences for helpfulness and safety. We utilised these models in their vanilla form, but also fine-tuned them on the curated training data of RegulaTome, employing the state-of-the-art QLora (Dettmers et.al. 2023) algorithm that makes adaptation of these foundation models to domain-specific tasks possible. The LLMs are instructed to produce structured output (JSON files), which renders any downstream processing and analysis tasks much easier. We use the recently released annotated RegulaTome corpus (Nastou et al., 2024) to evaluate the performance of RE against an established ground truth of PPI and GRN relations (for more details, see below). Generally, we report Precision, Recall, and F1-score values to estimate how well the model identifies and classifies PPI and GRN relationships specifically.

### 2.2. Evaluation Dataset

To assess the extraction of PPI and GRN relations, we utilised RegulaTome, a recently established resource (Nastou et al., 2024). This resource is, to our knowledge, the largest biomedical corpus for relation extraction (RE) available, comprising 2,521 scientific documents and 16,962 annotated relations. The RegulaTome corpus includes annotations for various types of relations, categorised based on their sign and directionality. It supports document-level annotations and facilitates the classification of relations by type. Among the 43 distinct relation types reported, we categorised 35 as PPIs and 2 as GRN relations (Supplementary Text 1). Upon annotation based on categories, we get 6,435 PPIs (Train:3870, Test: 1264 and Devel: 1301) and 1,076 GRNs (Train: 668, Test: 197, and Devel: 211).

The authors of the RegulaTome corpus (Nastou et al. 2024) trained a small transformer-based model (https://github.com/farmeh/RegulaTome_extraction) and reported an overall F1-score of 66.6% for their RE tasks as well as an F1-score of 91% for inter-annotator agreement (IAA) for all relationship types. In RegulaTome, relations were predicted for all possible pairs of detected entities, and if the entities were too far away to fit into the context window (128 tokens), they were discarded. Additionally, evaluation metrics (Precision, Recall and F1-Scores) were also reported individually for each type of relation. To match our definition of PPI and GRN RE tasks, we computed weighted average evaluation scores based on the assigned categories (Supplementary Text 1). This resulted in an overall F1-score of 73% for PPI relations (Precision of 74 % and Recall of 73%) and an F1-score of 58% (Precision of 59% and Recall of 57%) for GRN relations.

### 2.2. Base-prompt design

We have devised and applied a conversational prompt approach which extracts and then refines interactions between proteins and relations in gene regulatory networks. The prompt consists of three conversational steps. Breaking the process into steps ensured a systemic evaluation of precision and recall, as well as aligning the extraction to specific categories of relations. Additionally, a conversational multi-step approach would allow us to categorise the extracted relations based on their confidence level - the later the step in which a relation keeps getting reported, the higher its confidence.

For the implementation backend, we employed the langchain (langchain.com) framework that provides convenient wrappers to both communicate with local and API-deployed models. To ensure structured, parsable output from our system, we made use of the “function calling” capabilities of the latest model iterations. This option forces the model to fill a given pseudo-API reflecting the desired output structure - in our case, a JSON collection of directed triples of the form “[entity1, modifies, entity2]”.

### 2.3. Evaluation Types

We have performed PPI and GRN RE by using the two open-source LLMs as described in the previous section: Llama-8b and Llama-70b. For PPIs we have performed, the directionality of the relation was not accounted while for GRNs, the relations were evaluated by taking into account the directionality of the TF-to-Target relations. In both cases, the relations were extracted after each step of our multi-step prompt design. Additionally, we have relied on the original train-test split from RegulaTome in order to fine-tune Llama-8b and Llama-70b models to the training set of scientific texts. This allowed us to evaluate the Llama-8b and Llama-70b models over the test set in two different modes: *Fine-Tuned* and base pre-trained model (*Vanilla*). Finally, we also perform evaluations by either supplementing or not supplementing our prompts with examples of relations. In the case when examples are provided, we either: **i)** give examples of sentences which contain relations that need to be extracted (*Positive Examples*); **ii)** provide examples of sentences which contain relations which should be extracted (*Negative Examples*, i.e. we tell the LLM to not confuse associations or co-expressions with direct relations); **iii)** supplement both *Positive* and *Negative* Examples (from hereon referred to as *All Examples*). The examples that we provide have been tailored specifically for the type of relation that we aim to extract (i.e. PPI or GRN), and most importantly, they have been obtained from scientific texts which were not annotated in RegulaTome (Supplementary Text 2 and Supplementary Text 3). Additionally, we have also noticed that in any of the LLM configurations, there was no guarantee that the entity names in the extracted relations would match exactly with how they were annotated in RegulaTome. To address this issue, we additionally apply a prompt devised to extract 10 synonyms of each of the entities involved in the extracted relations. The prompt used for obtaining synonym names of the relation entities is: *“Generate at least 10 synonyms, genereric names or abbreviatons for the protein, gene or symbol provided by the user down below*.*”*. Such an approach was followed in order to mitigate effects such as when names of the entities in the extracted relations would not perfectly match the entities of the RegulaTome annotated relations, even though, according to our observations, it was clear that they were referring to the same entities. One such typical example is from the text *10666337* in the RegulaTome corpus where we had the following sentence: *“Human topoisomerase IIalpha and IIbeta interact with the C-terminal region of p53*.*”*. While in the RegulaTome corpus, the ‘*Human topoisomerase IIalpha→p53’* and *‘IIbeta→p53’* were annotated, the LLMs would instead extract ‘*Human topoisomerase IIalpha→p53’* and ‘*Human topoisomerase IIbeta→p53’*. This meant that only one of the two extracted relations would have been considered as a true positive if we only considered fixed names for the matching, even though it is clear that both relations should be considered as correctly extracted. In the said example, upon applying the synonym identification step, one of the synonyms of the *‘Human topoisomerase IIbeta’* was extracted to be *‘IIbeta’*, which then helped fixing the issue of the fixed matches. In order to assess the effect of introducing synonyms, evaluations *With* and *Without Synonyms* were then also performed for each LLM and prompt design configuration setting.

## 3. Results

### 3.1. PPI Evaluations from RegulaTome

As mentioned before, we evaluate the performance on the test set of the RegulaTome corpus using Precision, Recall and F1-score metrics. Here are provided the evaluation metrics for PPIs at each step of the prompt design for all combinations of: ***i)*** LLM model type: *Llama-8b* and *Llama-70b*; ***ii)*** in either *Vanilla* or *Fine-Tuned* mode; ***iii)*** with or without providing synonyms of the entities; and ***iv)*** when *Positive, Negative, All* and *No* examples are supplemented to the prompts. Evaluation results at Step-1 only, have been provided in Figure 2 (*Llama-8b* evaluations) and Figure 3 (*Llama-70b* evaluations) below.

**Figure 2.**
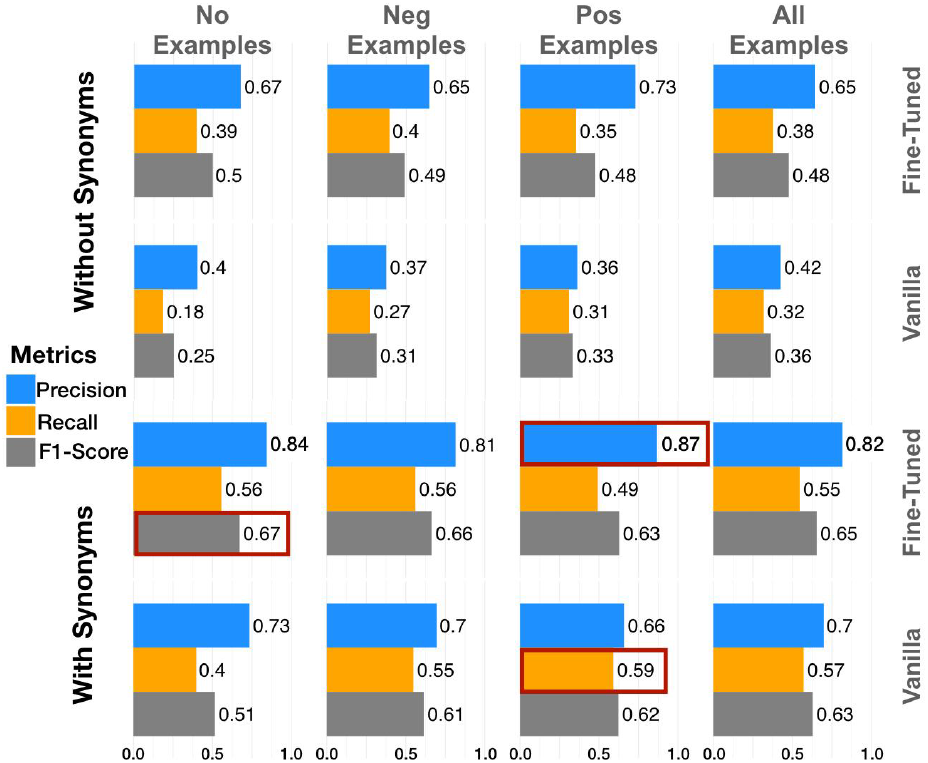
PPI RE evaluations at Step-1 using Llama-8b. Evaluations from Llama-8b in the Vanilla and fine-tuned styles without providing synonyms for entities in the extracted relations (top panel), as well as upon providing a list of synonyms for entities (bottom panel). In red are highlighted the top-performing models.

**Figure 3.**
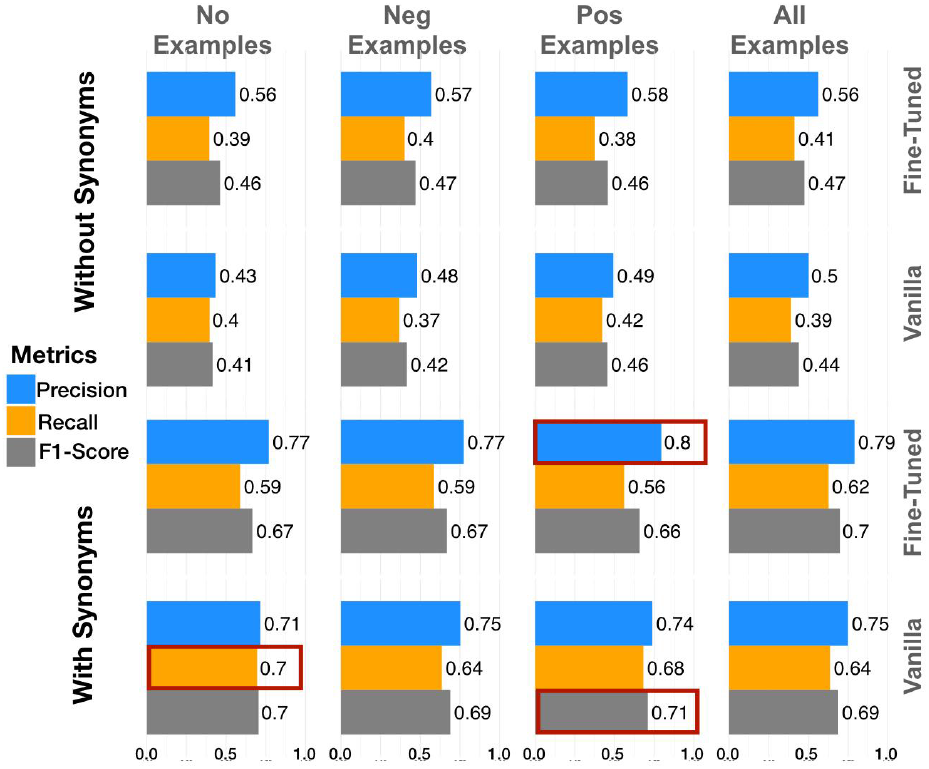
PPI RE evaluations at Step-1 using Llama-70b. Evaluations from Llama-70b in the Vanilla and fine-tuned styles without providing synonyms for entities in the extracted relations (top panel) as well as upon providing a list of synonyms for entities (bottom panel). In red are highlighted the top-performing models.

The evaluation results at Step-1 reveal that Llama-70b models perform consistently better than the Llama-8b counterparts (maximal Recall: 70% vs 59% and F1: 71% vs 67%) except for maximal Precision (80% vs 87%). As for evaluations with Llama-8b specifically, the fine-tuned models outperform their vanilla counterparts across all types of evaluations in terms of precision, recall and F1-score. This trend, however, disappears for the Llama-70b model, where we notice a slight degradation in overall F1-scores, mainly due to lower recall values. Precision, on the other hand, appears to be increasing in Fine-Tuned LLaMA-70b evaluations, while introducing lexical diversity of gene names through synonyms enhances the recall as well as its stability. Among the models tested, the *Llama-70b_FineTuned_AllExamples* and *Llama-8b_FineTuned_PosExamples* configurations deliver the highest precision, though with a noticeable trade-off in recall as steps progress, suggesting stricter but less comprehensive predictions. The *Llama70bV3_FineTuned_NegExamples* variant follows a similar pattern with slightly lower overall performance. The *Llama70bV3_Vanilla_PosExamples* variant, on the other hand, seems to be the best-performing model in terms of F1-score. Overall, the best evaluation metrics that we get (in Step-1) are: 87% precision (*Llama8bV1_FineTuned_PosExamples*), 70% recall (*Llama70bV3_Vanilla_NoExamples*) and an F1-score of 71% (*Llama70bV3_Vanilla_PosExamples*). Complete evaluation values across each step have been provided in Supplementary Table 1. Generally, we notice that every additional prompting step leads to an increase in precision, but at the expense of a lower recall, which leads to lower F1-scores as well.

### 3.2. GRN Evaluations from RegulaTome

Similar to the previous section, we present evaluation metrics for GRN at each step of the prompt design for all combinations of: ***i)*** LLM model type: *Llama-8b* and *Llama-70b*; ***ii)*** in either *Vanilla* or *Fine-Tuned* mode; ***iii)*** with or without providing synonyms of the entities; and ***iv)*** when *Positive, Negative, All* and *No* examples are supplemented to the prompts. Results have been provided in Figure 3 below.

**Figure 4.**
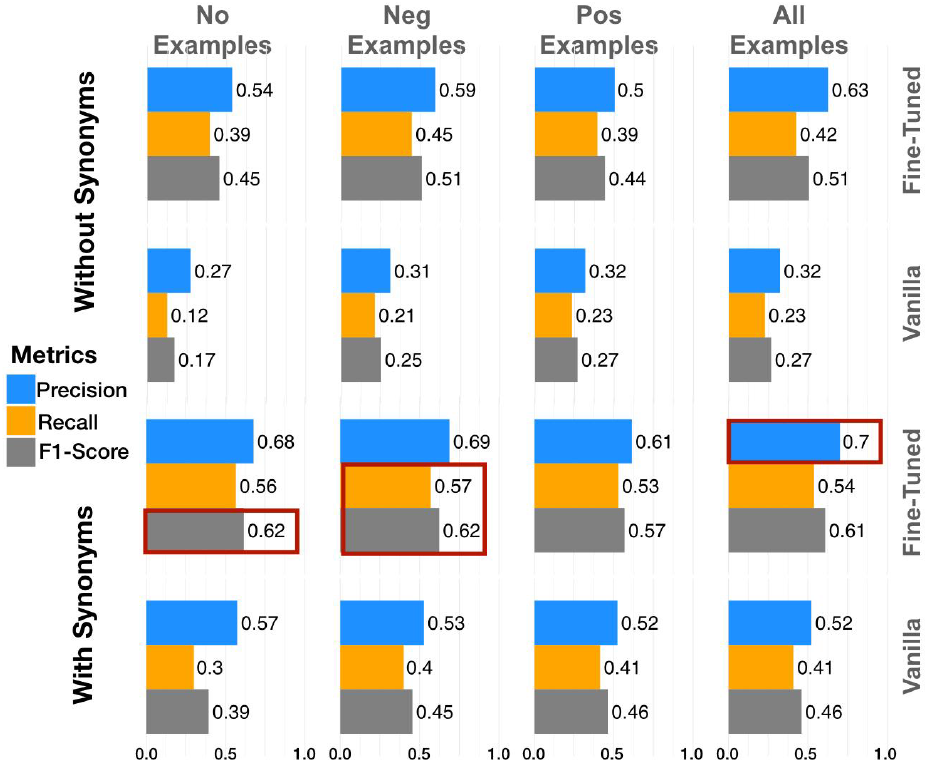
GRN RE evaluations at Step-1 using Llama-8b. Evaluations from Llama-8b in the Vanilla and fine-tuned styles without providing synonyms for entities in the extracted relations (top panel), as well as upon providing a list of synonyms for entities (bottom panel). In red are highlighted the top-performing models.

**Figure 5.**
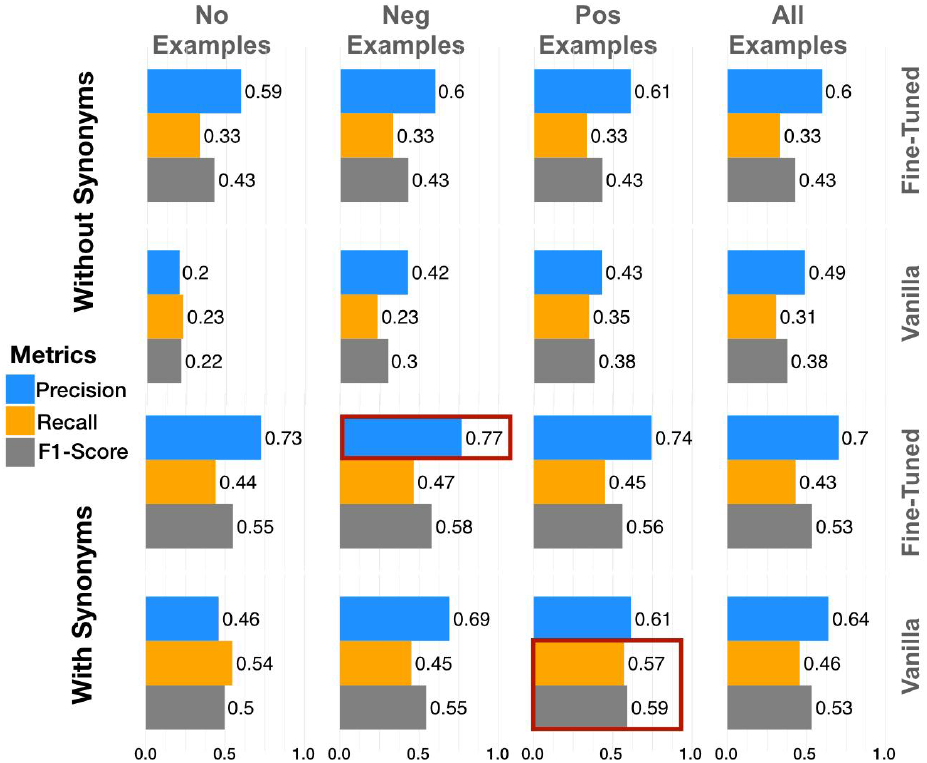
GRN RE evaluations at Step-1 using Llama-70b. Evaluations from Llama-70b in the Vanilla and fine-tuned styles without providing synonyms for entities in the extracted relations (top panel), as well as upon providing a list of synonyms for entities (bottom panel). In red are highlighted the top-performing models.

Generally, we observe overall better evaluation metrics in the Large LLaMA-70b model as compared to LLaMA-8b for the *Vanilla* mode of analysis. For the Fine-Tuned version of the models, on the other hand, this trend is reversed, and we have LLaMA-8b outperforming the larger LLaMA-70b model. Also, here we observe that fine-tuning of LLaMA-8b models seems to yield noticeable improvements in terms of precision, recall and F1-score values, while for LLaMA-70b models, the fine-tuned version improves on precision but loses recall (with F1-score remaining more or less similar). GRN REs seem to perform worse when compared to PPI REs, suggesting that LLMs might have more difficulty in extracting relations that consist of TF-Gene expression pairs as compared to interacting protein pairs. Overall, while these models are effective at making correct predictions when they do identify interactions (with a maximal precision of 77%), they miss a notable portion of relevant instances (maximal recall of only 57%), highlighting a precision-recall trade-off that becomes more pronounced over each step (see Supplementary Tables 1 & 2). The best evaluation metrics that we get (in Step-1) for GRN evaluations without prior knowledge of entities are: 77% precision (*LLaMA-70b_FineTuned_NegExamples*), 57% recall (*LLaMA-8b_FineTuned_NegExamples*, and *LLaMA-70b_Vanilla_PosExamples*) and an F1-Score of 62% (*LLaMA-8b_FineTuned_NegExamples*) – all when accounting for entity synonyms. Complete evaluation values across each step have been provided in Supplementary Table 2. There, similar to PPIs, we also notice that across each step, we have a general tendency for increased Precision and decreased Recall and F1-score across each step.

## 4. Discussion and Conclusions

Herein, we explored the capabilities of open-source LLMs, LLaMA-8b and LLaMA-70b, in extracting biological relations from scientific literature, focusing specifically on PPI protein interactions and TF-Gene relations relevant to GRNs. While LLMs are appealing for automatic mining of context-specific molecular interactions from literature, our findings suggest that models generally perform better in extracting PPIs (maximal Precision=87%, Recall=70% and F1=71%) compared to GRNs (maximal Precision=77%, Recall=57% and F1=62%). In terms of F1-scores, our best-performing models appear to be: *Llama70b_Vanilla_PosExamples_WithSynonyms* (Precision=74%, Recall=68% and F1=71%) for PPIs and *Llama8b_FineTuned_NoExamples_WithSynonyms* (Precision=68%, Recall=57% and F1=62%). The observations that LLMs tend to perform better in extracting PPIs as compared to GRNs (average differences PPI vs GRN: Precision=7–29%, Recall=-1–15% and F1=5–12%). Such differences seem to be in line with the evaluations reported in RegulaTome. Upon estimating weighted averages (based on size) of the evaluations reported in RegulaTome for each of the relation categories described in *Supplementary Text 1*, it resulted in an overall F1-score of 73% for PPI relations (Precision of 74 % and Recall of 73%) and an F1-score of 58% (Precision of 59% and Recall of 57%) for GRN relations. However, there are key differences in our approach. In our setup/experiment design, the LLM gets no prior information about the entities in the text and their location within the text. This is drastically different to the RegulaTome model, which provides only one pair of entities together with their position and the inter-spanning tokens. Additionally, our evaluations fall slightly short of the accuracy levels demonstrated by human annotation with an overall reported IAA F1-score of 91%, which suggests that models are not yet achieving consistent and reliable performance. In our case, the IAA of the LLMs would vary between 38% to 85%.

Key advantages of our prompt-based workflow include the use of open-source models and that we address the intrinsic limitations of annotation-based evaluations. Specifically, we are able to match variants of entities of the extracted relations through the use of synonyms without having the need to know a priori the names of the entities involved in the relations or their position within the text. For example, in (Chang et. al.), the authors evaluated prompts to extract PPIs from 6 different datasets by using proprietary models such as OpenAI’s ChatGPT (v3.5/v4) and Google’s Gemini. In this study, the most successful prompt achieved an F1 score on the complex datasets similar to our best performing scores (70%), however, in their case, the Genes/Proteins were tagged and numbered, meaning that they already provided the exact names of the entities that are known to be participating in the annotated relations. Similarly, (Rehana et al.) tested a range of fine-tuned BERT models against proprietary GPT-4. Their findings on a curated subset of the Human Protein Reference Database show supreme performance for the encoder-only models. Only with elaborate prompt engineering and temperature grid search, GPT-4 could close the gap when prompted on a sentence-level basis. (Degnan et al. 2024) extend on this view by not only comparing statistical methods, encoder-only and LLM models on the task of PPI extraction on the BioRED dataset (Luo et al. 2022), but also assessing graph-structured and true positive rates generated on a custom PubMed corpus in comparison to its corresponding UniProt standard. For gene-gene extraction, results on BioRED were generated on a sentence-level basis, combined with guiding LLMs by mentioning the ground truth proteins, making them competitive within the otherwise mixed results. The graphical analysis revealed that all three types of algorithms exhibited different network structures in terms of the chosen graph metrics, but the general low performance with regard to true positive relations doesn’t allow further conclusions. While BioRED is an extensively used annotated resource of biological entity relations, we opted not to include it in our evaluations. The primary reason for this is that our workflow is multi-task based, meaning that we are focused on extracting PPIs and GRNs specifically. The gene–gene relations in BioRED, however, are broadly defined and not limited to any single biological mechanism, as they can include various types of molecular relationships. The RegulaTome corpus, on the other hand, annotates types of relations which can be clearly categorised as either PPI or GRN. Another reason why we did not focus on BioRED is that they also annotated as relations indirect associations such as co-expression, functional associations, correlated activity, etc. In our workflow, we are instead explicitly interested in extracting direct relations between molecular entities.

Other advantages that this study offers are that we focus on extracting two different types of relations between biological entities, namely non-directional PPIs and directional GRNs (TF-to-Target), thus providing a comprehensive insight into molecular networks at different layers of the interactome. Additionally, in our workflow, the extracted relations are returned as structured JSON, thus facilitating downstream analysis and integration.

A limitation of our study is rooted in the RegulaTome corpus, which consists of short abstracts and paragraphs of scientific text (average number of tokens of 333, 336 and 325 for the train, dev and test sets in RegulaTome). In our view, performance on short texts cannot be easily extrapolated to entire manuscripts, which in principle are accessible to our approach. One obvious solution would be performing human annotation of a corpus of entire manuscripts. While this represents the most straightforward and intuitive approach, the annotation of thousands of manuscripts from multiple annotators all at once can be a daunting task.

Our work highlights the current potential of LLMs to aid in extracting and understanding complex biological interactions. When strictly looking at the best performing evaluation metrics of the RoBERTa-based model from RegulaTome, we achieve comparable numbers for GRNs (RoBERTa: Precision=59%, Recall=57% and F1=58% vs Llama: Precision=77%, Recall=57% and F1=62%) and PPIs (RoBERTa: Precision=74%, Recall=73% and F1=73% vs Llama: Precision=87%,

Recall=70% and F1=71%). However, we have to keep in mind that the RoBERTa-based models that were presented in the original RegulaTome study relied on highly informative pre-annotations (model inputs held both the entity type and their exact position in the tokenised text). In our workflow, on the other hand, we have no prior information about relation entities as well as their location and positional span. Additionally, our step-wise approach made it possible to classify the extracted relations based on their confidence: relations appearing across all the steps are to be classified as more robust and high-confidence; however, this comes at the expense of Recall. Also, our unique approach in introducing Synonyms of entities (generic names or abbreviations for the protein, gene or symbol extracted) seemed to mitigate many of the issues that are related to the correct evaluation of annotated text-based corpora. We believe that this work will pave the way for the development of improved models and base-prompt designs that can be used for the purpose of extracting molecular relations across domain-specific manuscripts.

## Availability

Codes with scripts and results have been provided in: https://github.com/dieterich-lab/LLM_Relations.

