## Supplementary Text for "Reading papers: Extraction of molecular interaction networks with large language models"

Supplementary Materials:

Supplementary Text 1 2
Supplementary Text 2 3
Supplementary Text 3 4

### Supplementary Text 1 - RegulaTome Relation Annotation

In RegulaTome we counted 12494 entities mentioned as well as 14842 total annotated relations (6398 categorised as PPIs and 6431 categorised as GRN relations) among 7503 entities.

The relations have been categorised as follows:

| PPI's | GRN's |
|---|---|
| Complex Formation, Catalysis Of Posttranslational Modification, Catalysis Of Ubiquitination, Catalysis Of Methylation, Catalysis Of Deacetylation, Catalysis Of Phosphorylation, Catalysis Of Dephosphorylation, Catalysis Of Acetylation, Catalysis Of Glycosylation, Catalysis Of Acylation, Catalysis Of Deneddylation, Catalysis Of Small Protein Conjugation, Catalysis Of Sumoylation, Catalysis Of Other Small Molecule Conjugation Or Removal, Catalysis Of Demethylation, Other Catalysis Of Small Molecule Conjugation, Catalysis Of Adp-Ribosylation, Catalysis Of Palmitoylation, Catalysis Of Neddylation, Catalysis Of Phosphoryl Group Conjugation Or Removal, Catalysis Of Deubiquitination, Catalysis Of Small Protein Conjugation Or Removal, Other Catalysis Of Small Protein Conjugation, Other Catalysis Of Small Protein Removal, Catalysis Of Geranylgeranylation, Catalysis Of Farnesylation, Catalysis Of Lipidation, Catalysis Of Prenylation, Catalysis Of Small Protein Removal, Catalysis Of Deglycosylation, Catalysis Of Desumoylation, Catalysis Of Depalmitoylation, Catalysis Of Deacylation, Catalysis Of Small Molecule Removal, Other Catalysis Of Small Molecule Removal | Regulation of gene expression, Regulation of transcription |

### Supplementary Text 2 - PPI Relation Extraction: Prompt and Examples

The prompt used to extract PPIs:

**Step 1:** *"You are a top-tier molecular biologist specialised in the field of molecular biology. Your task is to identify all the protein-protein interactions (PPIs) in the text, focusing on proteins involved in signalling pathways."*

**Step 2:** *"Review the above PPIs to determine whether they are specific to signalling pathways. Retain only signalling pathway interactions and remove the rest."*

**Step 3:** *"Review one more time the above PPIs to determine whether there are in the list relations that are of gene regulatory nature. Retain only those interactions that are PPIs and specific to cell signalling and remove those relations involving transcription factors to their gene targets."*

Below are provided *Positive* and *Negative* PPI relation extraction examples that supplement the prompts and help to guide the correct extraction of relations.

#### 1. Positive Examples

- *"This cytokine induces the p53 into a mutant-like conformation that forms a complex with Sp1"*: **p53 → Sp1**
- *"These findings suggest that the STAT3-NRF2 complex accelerates BLBC growth and progression by augmenting IL-23A expression."*: **STAT3 → NRF2**
- *"HIF1A forms a transcriptional complex with ARNT under hypoxia."*: **HIF1A → ARNT**.
- *"PRMT1 methylates cGAS and suppresses cGAS/STING signaling in cancer cellsPRMT1 methylates cGAS and suppresses cGAS/STING signaling in cancer cells"*: **PRMT1 → cGAS**
- *"TRAF6 ubiquitinates TGFβ type I receptor to promote its cleavage and nuclear translocation in cancer."*: **TRAF6 → TGFβ**
- *"AKT1 phosphorylates AKT1S1 at Thr-246."*: **AKT1 → AKT1S1**
- *"PIAS1 sumoylates PNKP in cells."*: **PIAS1 → PNKP**
- *"CBP, but not p/CAF, acetylates GATA-1 at two highly conserved lysine-rich motifs present at the C-terminal tails of both zinc fingers."*: **CBP → GATA-1**

#### 2. Negative Examples

- *"KRAS and BRAF cooperate in the MAPK signaling cascade to promote cell proliferation."* – Although the two proteins are in the same signalling system, the text does not provide evidence of a direct interaction.
- *"p53 and Protein MYC are both found in the same signaling complex."* – Incorrect assumptions based on co-occurrence or proximity.

- *“TNF and IL6 accumulate at DNA damage sites.”* – Co-localization but no evidence of direct relation/interaction between the two.
- *“Gene TNF regulates the expression of Gene IL6”* – Misinterpretation of genetic or signalling pathways as protein interactions.
- *“Prmt5 shares 80% sequence identity with Protein Prmt7, which is known to bind BRAF.”* – Incorrect assumptions based on structural similarity.
- *“PTEN was pulled down in a co-IP assay with CDKN2A.”* – Incorrect interpretations of experimental methods.

### Supplementary Text 3 - GRN Relation Examples

The prompt used to extract PPIs:

**Step 1:** *“You are a top-tier molecular biologist specialised in the field of molecular biology. Your task is to identify all the transcription factor (TF) to their gene target relations in the text that are involved in gene regulatory networks (GRNs).”*

**Step 2:** *“Review the above TF-to-Gene Target relations to determine whether they are specific to GRNs. Retain only GRN relations and remove the rest.”*

**Step 3:** *“Review one more time the above TF-to-Gene Target relations to determine whether there are in the list relations that are of protein-protein interaction nature. Retain only those interactions that are GRNs and remove those relations involving interaction between proteins.”*

Below are provided *Positive* and *Negative* GRN relation extraction examples to guide the correct extraction of relations.

#### 1. Positive Examples

- *“MYC target genes that are involved in cell cycle such as Cyclin D1”*: **MYC → Cyclin D1**
- *“STAT3 can induce the expression of anti-apoptotic genes like Bcl-2, which help in cell survival.”*: **STAT3 → Bcl-2**
- *“Tbx1 activates transcription of the fibroblast growth factor genes Fgf8 and Fgf10 to maintain proliferative expansion and inhibit differentiation of cardiopharyngeal precursor cells”*: **Tbx1 → Fgf8** and **Tbx1 → Fgf10**
- *“Overexpression of MECP2 leads to the suppression of IFN-γ transcription, which is linked to impaired TH1 responses in both children and mice with MECP2 duplication syndrome.”*: **MECP2 → IFN-γ**
- *“In humans, FOXO regulates the expression of core small RNA pathway genes, including AGO2.”*: **FOXO → AGO2**.

#### 2. Negative Examples

- *“This cytokine induces the p53 into a mutant-like conformation that forms a complex with Sp1”* – This is complex formation and does not involve transcription factor to gene relations.
- *“HIF1A forms a transcriptional complex with ARNT under hypoxia.”* – This relation represents two transcription factor proteins interacting with each other, and the text does not reflect that they target the regulation of any specific gene.
- *“PRMT1 methylates cGAS and suppresses cGAS/STING signaling in cancer cells”* – This is a methylation interaction and not a transcription factor to gene relation.
- *“Gene MYC and gene STAT3 share a common promoter region.”* – This is not explicitly a relation between a transcription factor and its target gene.
- *“AKT1 phosphorylates AKT1S1 at Thr-246.”* – This is a phosphorylation interaction and not a transcription factor to gene relation.
